# “Flower power”: how flowering affects spectral diversity metrics and their relationship with plant diversity

**DOI:** 10.1101/2023.12.11.571067

**Authors:** Michela Perrone, Luisa Conti, Thomas Galland, Jan Komárek, Ondřej Lagner, Michele Torresani, Christian Rossi, Carlos P. Carmona, Francesco de Bello, Duccio Rocchini, Vítězslav Moudrý, Petra Šímová, Simonetta Bagella, Marco Malavasi

## Abstract

Biodiversity monitoring is constrained by cost- and labour-intensive field sampling methods. Increasing evidence suggests that remotely sensed spectral diversity (SD) is linked to plant diversity, holding promise for monitoring applications. However, studies testing such a relationship reported conflicting findings, especially in challenging ecosystems such as grasslands, due to their high temporal dynamism and variety. It follows that a thorough investigation of the key factors, such as the metrics applied (i.e., continuous, categorical) and phenology (e.g., flowering), influencing such a relationship is necessary. Thus, this study aims to assess the applicability of SD for plant diversity monitoring at the local scale by testing six different SD metrics while considering the effect of the presence of flowering on the relationship and resampling the original data to assess how spatial resolution affects the results. Taxonomic diversity was calculated based on data collected in 159 plots with 1.5 m ×1.5 m experimental mesic grassland communities. Spectral information was collected using a UAV-borne sensor measuring reflectance across six bands in the visible and near-infrared range at ∼2 cm spatial resolution. Our results show that, in the presence of flowering, the relationship is significant and positive only when SD is calculated using categorical metrics. Despite the observed significance, the variance explained by the models had very low values, with no evident differences when resampling spectral data to coarser pixel sizes. Such findings suggest that new insights into the possible confounding effects on the SD∼plant diversity in grassland communities are needed to use SD for monitoring purposes.

## 1 Introduction

The current unprecedented biodiversity change rate experienced worldwide (Wilting et al., 2017) makes species conservation one of the most pressing priorities of our times. While there has been a growing interest in species conservation, biodiversity evaluation and monitoring are still limited and lack standardized methods for quick and scalable data gathering (Palmer et al., 2002; Skidmore et al., 2015; Wang & Gamon, 2019). Among the available approaches for the remote monitoring of biodiversity, increasing evidence suggests that remotely sensed spectral diversity (SD) is linked to plant diversity (Schweiger et al., 2018). The SD concept was originally framed within the spectral variation hypothesis (SVH; Palmer et al., 2002), which assumes an indirect relationship between spectral and plant diversity through environmental ‘surrogacy’, i.e., the landscape heterogeneity captured by RS (Hauser et al., 2021; Torresani et al., 2019). When relying on medium-coarse spatial resolution data, the link between spectral and plant diversity can only be of indirect nature, as pixel size exceeds the size of individual plants. Conversely, the high spatial resolution of the multispectral data provided by uncrewed aerial vehicles (UAVs) holds promise for being able to capture the direct link between spectral and plant diversity at the leaf and canopy level in herbaceous communities (Rossi et al., 2021a; Thornley et al., 2023). However, when the pixel size reflects that of individuals (or even the sub-individual level), the intra-specific (or intra-individual) variability in the plant optical traits can play a role in the SD-biodiversity relationship.

Studies testing the relationship between SD and plant diversity at fine scales have reported conflicting findings, especially in challenging ecosystems such as grasslands, whose biodiversity is threatened by their depletion, degradation, and fragmentation due to anthropogenic factors (Thornley et al., 2023). Their vast global distribution (Gibson, 2009) and the broad range of ecosystem services they provide make grassland conservation a priority (Hein et al., 2006; Suding, 2011), yet their extensive monitoring is prevented by logistical constraints. Due to its efficiency and cost-effectiveness, SD-based plant diversity monitoring offers an appealing alternative to traditional sampling techniques in grasslands.

Nonetheless, the path towards the routine implementation of SD into grassland monitoring is hampered by the high temporal and spatial variability of such ecosystems, which often exhibit a complex community structure, particularly in natural or semi-natural grasslands (Wilson et al., 2012). Recent studies have highlighted the crucial role played by plant phenology (e.g., flowering, leaf emergence and senescence; Rossi et al., 2021b; Thornley et al., 2022) of spectral data when addressing the relationship between spectral diversity and plant diversity in grasslands. Despite the dynamism of grassland communities, acquiring repeated intra-annual spectral data is time- and resource-consuming. It follows that spectral data are usually acquired to ensure a temporal match with vegetation surveys when the presence of dead biomass and exposed soil is supposed to be the lowest (Asner, 1998). However, this also implies that the species present would potentially be flowering, causing unwanted additional sources of spectral variability. The presence of flowering is likely to have an impact on the SD of a community, as it leads to a high variation in important optical traits and, thus, in the spectral signatures of the different species (Conti et al., 2021; Fassnacht et al., 2022; Gholizadeh et al., 2019; Thornley et al., 2022). Being non-photosynthetic organs with distinct spectral features, especially in the visible domain (Schiefer et al., 2021), flowers will affect the calculation of SD, and the extent of this impact is likely to depend on the characteristics of the SD metrics applied.

The metrics used to quantify SD can be either continuous (i.e., based on variation in traditional vegetation reflectance indices or on the full spectral information) or categorical (i.e., based on the categorisation of the spectral space; Féret and Asner, 2014) (Wang and Gamon, 2019). Although continuous SD metrics are the most represented and widely tested, they show weakness in capturing plant diversity by measuring the degree of contrast in RS optical data (Fassnacht et al., 2022). In the case of few - but substantially distinct in their spectral signature - species present, the spectral heterogeneity can be very high even if the plant diversity is low (Fassnacht et al., 2022). Following the same reasoning, continuous SD metrics are likely to be influenced by confounding factors like flowering, as well as soil presence and amount of biomass. In contrast, categorical SD metrics are considered less sensitive to extreme reflectance values (e.g., from background material) as such values would represent distinct categories alongside other equally significant ones (Rossi et al., 2021a). Even though some studies have tested and compared different SD metrics (Frye et al., 2021; Gholizadeh et al., 2020, 2018; Perrone et al., 2023; Rossi et al., 2021a; Schmidtlein and Fassnacht, 2017), there is still no general consensus on which metrics would be the best proxy for plant diversity in the presence of flowering.

Here, building on top of the previous results presented by Conti et al. (2021), we further explore the relationship between spectral and plant diversity by relying on species and multispectral UAV data from experimental grassland communities located in a mesic meadow in South Bohemia (Czech Republic). In the present study, we aim to assess the applicability of SD for plant species richness monitoring at the local scale by testing six different SD metrics (detailed in Table 1) and considering flowering as a factor that potentially confounds the relationship. Moreover, we investigate if varying the spatial resolution (from 2 cm to 5 cm) influences the observed relationship.

**Table 1.**
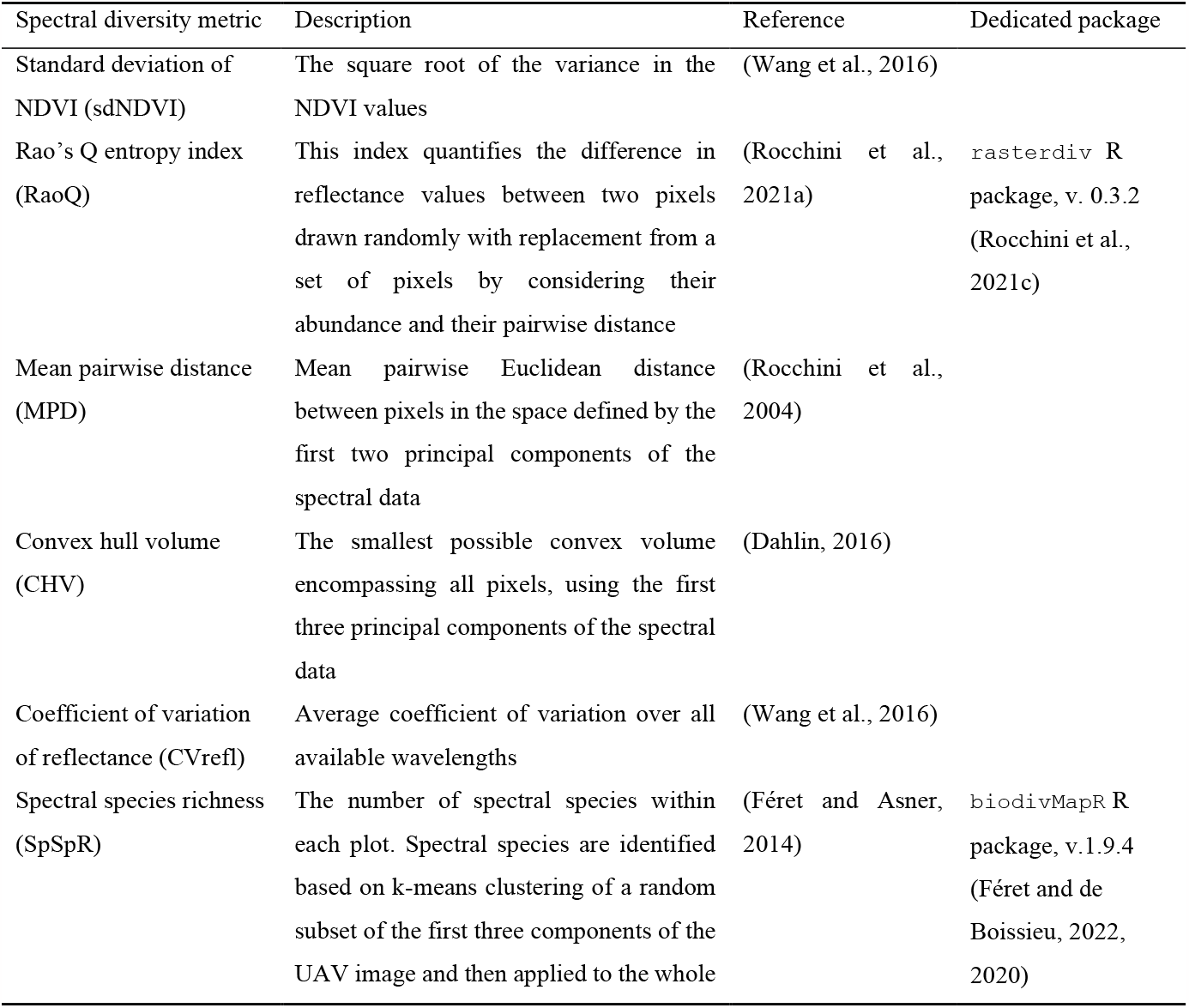
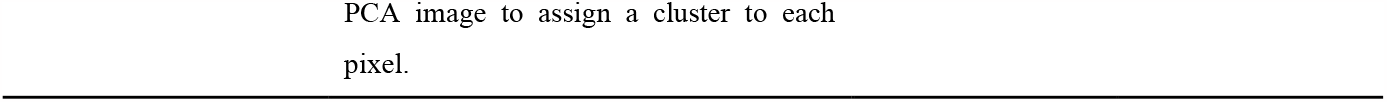
Description of the spectral diversity metrics used in this study.

| Spectral diversity metric | Description | Reference | Dedicated package |
| --- | --- | --- | --- |
| Standard deviation of NDVI (sdNDVI) | The square root of the variance in the NDVI values | (Wang et al., 2016) |  |
| Rao's Q entropy index (RaoQ) | This index quantifies the difference in reflectance values between two pixels drawn randomly with replacement from a set of pixels by considering their abundance and their pairwise distance | (Rocchini et al., 2021a) | `rasterdiv` R package, v. 0.3.2 (Rocchini et al., 2021c) |
| Mean pairwise distance (MPD) | Mean pairwise Euclidean distance between pixels in the space defined by the first two principal components of the spectral data | (Rocchini et al., 2004) |  |
| Convex hull volume (CHV) | The smallest possible convex volume encompassing all pixels, using the first three principal components of the spectral data | (Dahlin, 2016) |  |
| Coefficient of variation of reflectance (CV<sub>refl</sub>) | Average coefficient of variation over all available wavelengths | (Wang et al., 2016) |  |
| Spectral species richness (SpSpR) | The number of spectral species within each plot. Spectral species are identified based on $k$-means clustering of a random subset of the first three components of the UAV image and then applied to the whole | (Féret and Asner, 2014) | `biodivMapR` R package, v.1.9.4 (Féret and de Boissieu, 2022, 2020) |

## 2. Material and Methods

### 2.1 Field data

Our study is based on vegetation data collected in a permanent grassland experiment of the University of South Bohemia located in a mesic meadow in the Vysočina region (South Bohemia, Czech Republic, 49.331N, 15.003E). As Galland et al. (2019) described, the sowing experiment consists of 40 mesic grassland communities covering independent gradients of plant functional and phylogenetic diversity. These experimental communities derive from a specific treatment, i.e., either the sowing of a combination of 6 species drawn from a pool of 19 mesic meadow species naturally present in the area or from monoculture treatments. Each 6-species community was sown in two randomly situated 1.5 m × 1.5 m plots (one fertilised and one unfertilised), while monoculture plots were sown in three replicates each. Thus, the study site encompasses a total of 196 plots, with a buffer zone of 0.5 m between them (Figure 1). Due to the subsequent processing of UAV data, only 159 out of the 196 plots were considered in this study.

**Figure 1.**
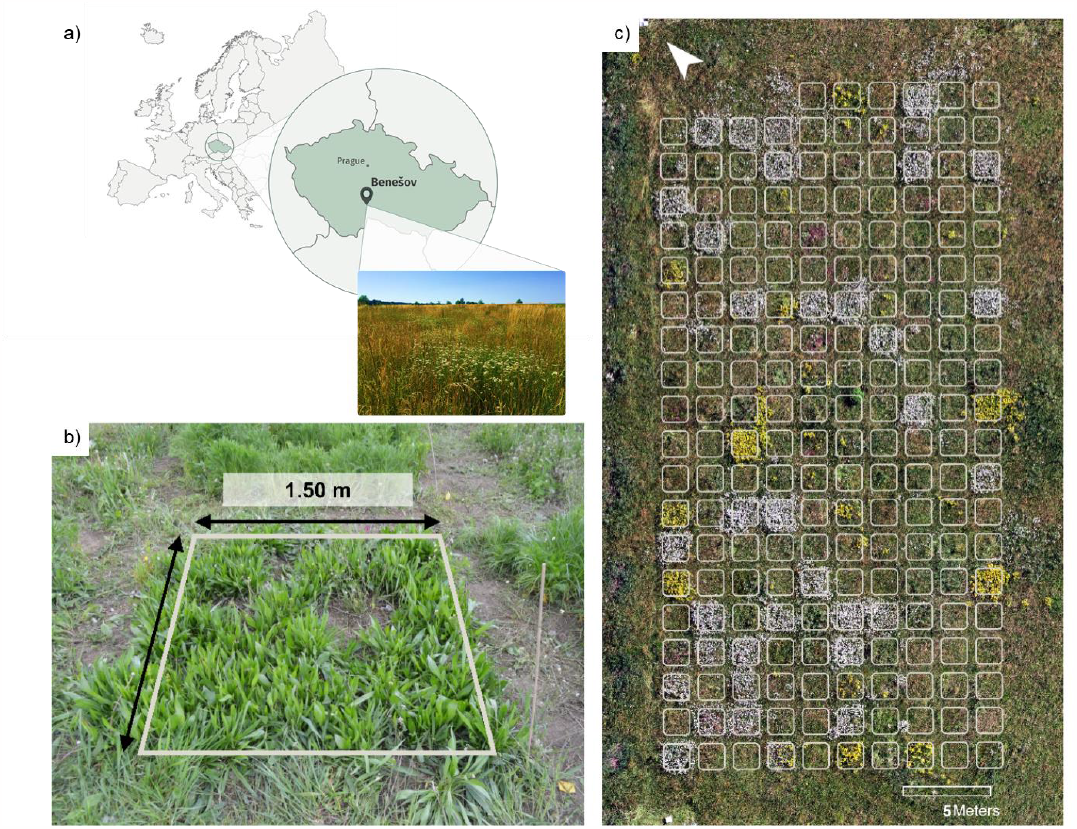
(a) Location of the study area. (b) Example of a 1.5 m × 1.5 m surveyed plot. (c) natural-colour mosaic of the whole study area.

Plant species composition within each plot was measured in May 2019. The actual species richness (SR) of the experimental communities varied between 12 and 36 species due to the later plot invasion by spontaneously colonising species.

### 2.2 Spectral data

We used multispectral UAV imagery with ∼2 cm spatial resolution (80% frontal and 70% side tile overlapping, 30 m flight altitude) acquired at the peak of the growing season (July 2019) using a Kingfisher multicopter (Robodrone industries, Brno, Czech Republic) equipped with a Micro-MCA6 (Tetracam Inc., Chatsworth, CA, United States) camera measuring reflectance across six 10-20nm-width bands in the 490 – 900 nm range of the electromagnetic spectrum. The spectral data obtained were processed using a Structure from Motion and Multi-View Stereo algorithms in the Metashape (Agisoft LLC, St. Petersburg, Russia) image-matching software to generate an ortho-mosaic. Georeferencing was performed with spatial error of 2 cm using ground control points surveyed through a Leica GPS1200 GNSS receiver (Leica Geosystems AG, Heerbrugg, Switzerland) in RTK mode. The ortho-mosaic was radiometrically calibrated using white and grey calibration targets for which the spectral properties were known through spectrometer measurements.

The ortho-mosaic portions exhibiting processing artefacts were excluded from further analyses, resulting in 37 out of 196 study plots being left out.

Previous studies have reported that the presence of bare soil, due to its substantially differing-from-vegetation spectrum, proved to significantly deteriorate SD performance in quantifying plant diversity (Gholizadeh et al., 2018; Hauser et al., 2021). Moreover, the spectral signal has proven to be sensitive to the vertical structure of the plant community, since a complex vertical structure of the community may generate an “occlusion effect” that would lead to the obscuration of shorter species and the presence of shadowed pixels (Conti et al., 2021). Here, we masked shadowed areas and non-vegetated pixels from the multiband ortho-mosaic to mitigate their influence on SD. Bare soil pixels were masked using an NDVI threshold of 0.3, while shadowed pixels were identified and removed through an Expectation Maximization (EM) unsupervised classification. Moreover, the presence of flowering plants within the plots was assessed visually from an RGB composite and attributed to each plot (i.e., a plot was classified as flowering given >10% coverage of flowering). Finally, we rescaled the ortho-mosaic to 5-cm spatial resolution (method: bilinear) to test if coarsening the spatial resolution can mitigate the effect of intraspecific (and intraindividual) variability of optical traits.

### 2.3 SD metrics

We calculated SD within the 1 m × 1 m core area of each 1.5 m × 1.5 m experimental plot by applying six different spectral diversity metrics, as there is no consensus on the best-performing method to quantify SD. Specifically, we computed six continuous SD metrics, namely, the standard deviation of NDVI (sdNDVI), the Rao’s Q entropy index (RaoQ), the mean pairwise distance of reflectance values (MPD), the coefficient of variation of reflectance of the full spectral range (CVrefl), and the convex hull volume (CHV). We also computed one categorical metric, i.e., spectral species richness (SpSpR) (please see Table 1 for further details on the metrics used). For the spectral species mapping we used the biodivMapR R package (for a complete description of the method, please see Féret and de Boissieu, 2020). To perform the mapping, the number of k clusters was set to 50, being a realistic number of the spectral species present based on ground data and as a trade-off between performance and computational efficiency. The unsupervised clustering of a random subset of pixels was repeated eight times and the SpSpR value assigned to each plot is the mean SpSpR value calculated based on such repetitions.

### 2.4 Statistical analyses

To test the relationship between SR and each of the metrics tested, we modelled the variation in SD through Generalized Linear Mixed Effects Models (Mcculloch and Neuhaus, 2014) (Gamma family, log link) using the lme4 R package *v*.*1*.*1-33* (Bates et al., 2015). In our models, SR and the flowering binary variable (presence/absence) were used as fixed-effect predictors. At the same time, the type of grassland community (i.e., monoculture, low/high functional diversity, low/high phylogenetic diversity, fertilised/unfertilised) was included as a random effect to account for the interdependence among communities in our dataset. To assess the goodness of fit for our models, we calculated for each model marginal (R^2^_m_) and conditional (R^2^_c_) pseudo-R squared (Nakagawa et al., 2017). These metrics quantify the proportion of variance explained by fixed effects and the complete model (i.e., both fixed and random effects), respectively. We calculated R^2^_m_ and R^2^_c_ using the r.squaredGLMM() function of the MuMIn R package, *v*.*1*.*43*.*17*. Data and scripts are provided at https://github.com/MichelaPerrone/SVH_Benesov.git under CC-BY license.

## 3. Results

At both the spatial resolutions tested, the models showed a significant and positive relationship between SR and SD only when SD was measured using categorical metrics (i.e., spectral species richness) (Table 2; Figure 2). Conversely, when the SD response variable was calculated using continuous metrics, only flowering showed a significant and positive association with SD (Table 2; Figure 2).

**Table 2.** Summary of the models’ results. Columns refer to the SD metric used as the response variable; rows refer to the coefficient estimates (coeff. est.) of the fixed-effect explanatory variables (log-transformed values) and the models’ marginal and conditional R2 values.

| Spatial resolution: 2 cm |  |  |  |  |  |  |
| --- | --- | --- | --- | --- | --- | --- |
|  | sdNDVI | RaoQ | MPD | CHV | CVrefl | SpSpR |
| SR coeff. est. | $-4.99\text{e}^{-3}$ | -0.02 | -0.02 | 0.06 | $-1.89\text{e}^{-5}$ | 0.02* |
| Flowering coeff. est. | 0.12*** | 0.34*** | 0.30*** | 0.65*** | 0.22*** | -0.02 |
| $R^2_m$ | 0.24 | 0.39 | 0.37 | 0.35 | 0.51 | 0.05 |
| $R^2_c$ | 0.29 | 0.50 | 0.48 | 0.44 | 0.58 | 0.13 |
| Spatial resolution: 5 cm |  |  |  |  |  |  |
|  | sdNDVI | RaoQ | MPD | CHV | CVrefl | SpSpR |
| SR coeff. est. | $-5.94\text{e}^{-3}$ | -0.02 | -0.02 | 0.05 | $4.66\text{e}^{-3}$ | 0.04** |
| Flowering coeff. est. | 0.15*** | 0.34*** | 0.32*** | 0.78*** | 0.21*** | $1.46\text{e}^{-3}$ |
| $R^2_m$ | 0.28 | 0.37 | 0.36 | 0.41 | 0.38 | 0.06 |
| $R^2_c$ | 0.29 | 0.49 | 0.47 | 0.48 | 0.50 | 0.16 |

**Figure 2.**
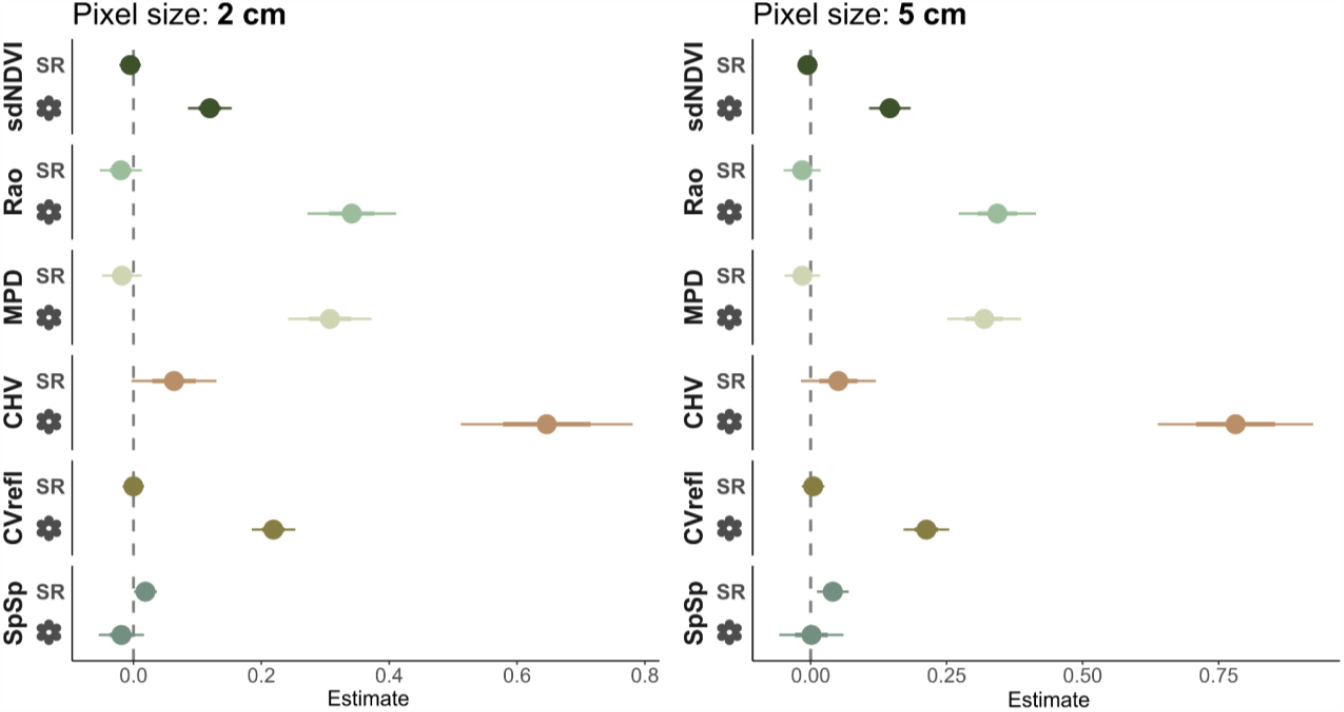
Coefficient plots of the mixed models illustrating the estimated (log-transformed) values for each explanatory variable, along with their corresponding 50% (inner error bars) and 95% (outer error bars) confidence intervals. Colours refer to the different SD metrics used as the response variable.

The influence of flowering plants on continuous metrics was substantial, with the variation in SD calculated through such metrics. This was reflected by the relatively high R^2^_m_ values obtained for models considering continuous SD metrics, ranging between 0.24 (sdNDVI, 2-cm spatial resolution) and 0.51 (CVrefl, 2-cm spatial resolution).

In contrast, models based on spectral species richness had very low R^2^_m_ values (Table 2; Figure 3), with no evident differences observed between the two spatial resolutions. Despite the significant relationship between SD and SR, the low variance explained by the fixed effects in the spectral species richness models suggests that other factors play a more prominent role at this ecological scale and within these settings.

**Figure 3.**
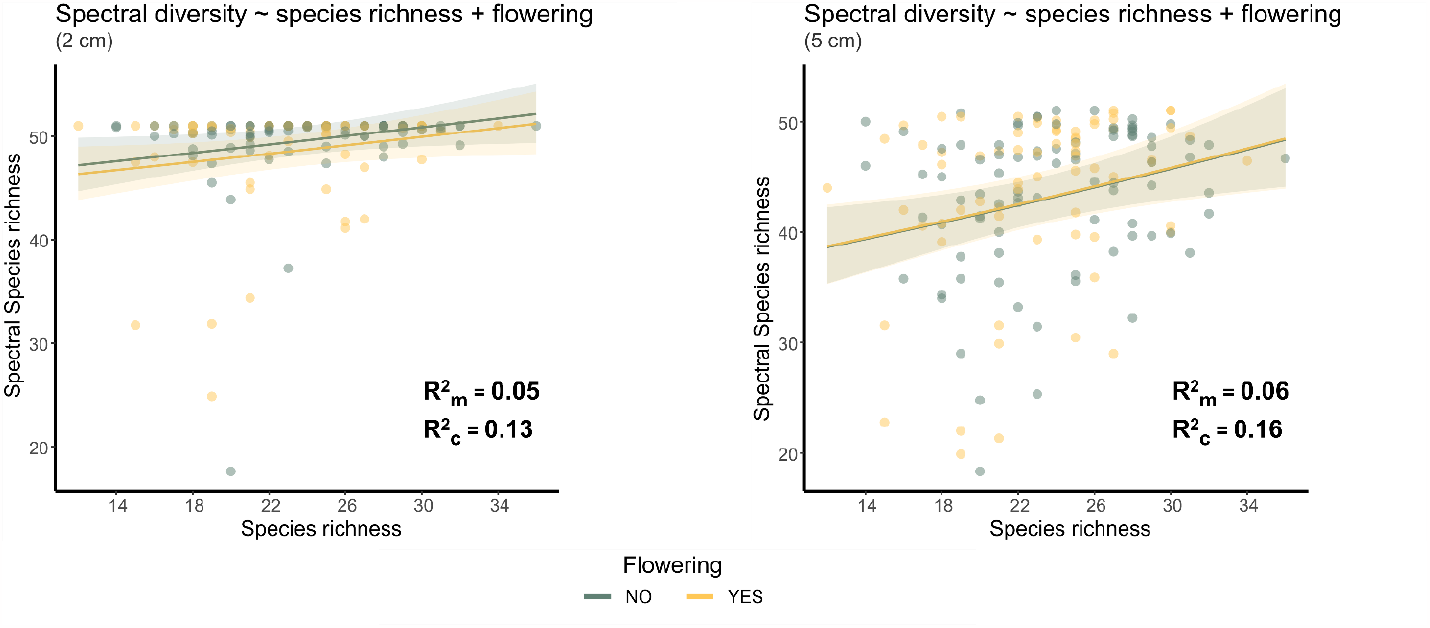
Predicted relationships between SD (calculated as spectral species richness) and species richness derived from the mixed-effect models for the two spatial resolutions tested based on the presence of flowering (see legend).

Moreover, we observed that a too-high spatial resolution, such as a 2-cm resolution, is not beneficial when using categorical metrics. Indeed, in such a case, we observed that a high number of spectral species are detected in most of the plots, regardless of the actual ground species richness (Figure 3).

## 4. Discussion

### 4.1 SD metrics

The community’s phenological pattern impacts the effective application of SD for biodiversity monitoring as plant species exhibit distinct physiological and structural characteristics at different phenological stages, causing changes in their spectral signatures. The presence of flowers markedly different in their spectral characteristics from plant green parts impacts the use of continuous SD metrics due to their sensitivity to extreme reflectance values. Notably, the range of spectral information considered, i.e., bands at specific wavelengths (sdNDVI) or all available spectral bands (RaoQ, MPD, CHV, CVrefl), did not improve substantially the performance of the continuous metrics, thus indicating that using only a subset of the available spectral information is not what limits the successful application of SD. To use any of the continuous SD metrics tested, the acquisition timing of spectral data turns out to be relevant. Indeed, an additional challenge to the applicability of the SVH for estimating biodiversity is the asynchrony in phenology among species present in the observed community, as the number of co-present phenological stages (both at the leaf- and flower-level) in a plot confounds the observed SD (Fassnacht et al., 2022; Thornley et al., 2022). Thus, the timing of data acquisition is paramount in discerning whether a connection between SD and species diversity can be observed. For this reason, categorical metrics are highly advantageous in this context, as they ideally enable overcoming the issue of seasonal change in plant optical properties due to species phenology. Indeed, this type of metric ideally allows the identification of spatial units with consistent spectral homogeneity through time (Perrone et al., 2023). Despite their advantages, the successful use of categorical metrics for estimating spectral diversity is challenged by the necessity of selecting the proper settings by the user, namely the optimal number of clusters into which the spectral space should be partitioned. While for hyperspectral data it is possible to use virtual dimensionality to get the number of clusters, in the case of multispectral data this requires going through a time-consuming trial-and-error procedure that should consider the specific plant communities observed, the reliability of the results, and the computation intensity (Féret and de Boissieu, 2020; Perrone et al., 2023; Rocchini et al., 2021b), it should also be noted the critical role played by the spatial resolution of the data used. Previous studies on SD highlighted how pixel size affects the relationship between spectral and plant diversity, which becomes considerably weaker with progressively coarser resolutions (Gholizadeh et al., 2018, 2019; Wang et al., 2016). However, the dependence of the relationship on pixel size is contingent on the SD metrics used (Gholizadeh et al., 2018) and sampling plot size, as well as on the specificity of the study site (Gholizadeh et al., 2022). Thus, we argue that when relying on very fine spatial resolutions that exceed the necessary scale for capturing individual objects, the risk is to introduce noise into the data (Conti et al., 2021), resulting in saturation in the number of clusters (i.e., spectral species) identified within each area unit (e.g., plot) (Figure 3). Therefore, users must strike a balance between spatial resolution and the desired level of spectral discrimination to avoid this issue and obtain meaningful results.

### 4.2 Improvements achieved with pixel masking

The complexity of the herbaceous communities’ vertical structure can lead to a negative relationship between SD and plant diversity due to the “occlusion effect” caused by taller species (Conti et al., 2021). Given such previous knowledge, we applied a shadow and bare soil masking on the original ortho-mosaics to improve the correlation and focus on the specific issues we wanted to test (i.e., the influence of flowering on SD). Indeed, as shown by Gholizadeh et al. (2018), filtering out pixels that capture information from various sources other than plants improves the performance of SD metrics, allowing to (partially) correct for such confounding factors. In principle, such correction is possible when relying on data with fine spatial resolution (e.g., proximal or UAV-borne sensors) since the spectral signature of single pixels is more likely to belong to a single object type. Moving to coarser spatial resolutions, such as with airborne and spaceborne sensors, spectral unmixing would be required to correctly identify mixed pixels and correct for bare soil presence, as it allows estimating the per-pixel percentage of end members (Asner and Heidebrecht, 2002; Gholizadeh et al., 2018; Rossi and Gholizadeh, 2023).

### 4.3 Additional confounding factors

In light of the low explanatory power of the models based on spectral species richness, additional confounding factors may have played a role in the computed SD. As highlighted by Rossi et al. (2021a), the presence and ratio of live and dead biomass, together with the total biomass (Villoslada et al., 2020), may noticeably influence the spectral signal, leading to higher spectral variability. Yet, the extent of such an impact is dependent upon the type of ecosystem (Rossi et al., 2021a), as well as the specific characteristics of the plant community observed (e.g., specific functional and morphological features), particularly at high spatial resolutions (Villoslada et al., 2020), supporting the context dependency of the relationship between spectral and plant diversity (Fassnacht et al., 2022). Other factors that have potentially affected our results are the number of species present and the composition of the experimental communities. Indeed, it has been previously observed that the species’ life forms (e.g., graminoids, forbs and legumes) and their prevalence in the community affect optical SD metrics (Gholizadeh et al., 2019; Wang et al., 2018b; Rossi et al., 2021a). Moreover, Imran et al. (2021) reported that the SD-plant diversity link holds better in artificial, species-poor ecosystems, while it is weaker in species-rich natural grasslands. Despite originating from an experiment, our study site has undergone a (partial) re-naturalisation due to the spontaneous colonisation by local species, thus increasing the species richness observed within each plot. Such a decrease in strength of the SD-plant diversity link in high species richness habitats further hinders the operational applicability of SD for biodiversity monitoring purposes in natural and semi-natural habitats.

### 4.4 Limitations

In the present study, we were interested in assessing if the simple presence of flowering has a confounding effect on the relationship between plant and spectral diversity measured through different metric types. Therefore, flowering was included in the analyses as a binary (presence/absence) variable. We did not attempt a qualitative or quantitative estimation of the flowering presence, as it would have diverted from our original aim. While we acknowledge that gathering information on the specific flower colour, size, and spatial cover would help characterise the presence of flowering within each plot, such types of data would not have been possible to obtain within our study settings (both in terms of ground and spectral data). Nevertheless, it is a promising topic for future research, which could be viable using hyperspectral data and ground data on flowering plants, and that would help get further insight into the confounding effect that flowering exerts on SD.

Moreover, the experimental setup on which our study is based would have determined the impossibility of further testing the influence of spatial resolution. Indeed, due to the small core plot size (1 m × 1 m) and pixel masking to reduce the impact of bare soils and shadowed areas, further rescaling (i.e., over a 5-cm spatial resolution) would have resulted in few pixels available for SD computation and, thus, introduced a major source of error (i.e., small sample size). Besides, our aim was to assess the direct link between SD and plant diversity, which implies the matching between pixel and individual size. Thus, further coarsening the spatial resolution would have been in contrast with our assumptions.

Additionally, we did not consider abundance-based indices (e.g., Shannon’s H) for both plant and spectral diversity due to the possible mismatch between the actual field-sampled species abundance and the retained spectral information after pixel masking. However, while species richness is the most widespread species diversity metric in SD studies, we acknowledge that abundance-based metrics have proven to be more strongly related to SD on several occasions (Oldeland et al., 2010; Torresani et al., 2019; Wang et al., 2018).

Finally, due to the unavailability of total and relative (live, dead) biomass data, as well as data on life forms abundance, we could not take into account such confounding factors in our study. To gain a deeper understanding of how all the confounding factors identified so far (i.e., bare soil, phenological features, biomass, vertical complexity, community composition) and their potential interactions affect the plant-spectral diversity relationship, future experiments should be designed to consider them simultaneously and assess their impact on different types of grasslands.

## 5. Conclusions

The reliability of using SD to monitor plant diversity is a matter of controversy and may need more consistency in specific settings, especially in dynamic ecosystems such as grasslands. In this study, we investigated the SD-biodiversity relationship in mesic grassland communities by testing how flowering presence may influence the estimation of plant species richness using different SD metrics. According to our results, the presence of flowering proved to impair the ability of continuous SD metrics to reflect plant diversity, with flowering likely being the main source of spectral variance within plots. On the contrary, categorical SD metrics appear less influenced, confirming the better suitability of this type of metrics observed in previous studies (Rossi et al., 2021a). Nevertheless, when calculating SD using categorical metrics, species richness only explains a small portion of the variability in spectral heterogeneity at both spatial resolutions tested. We hypothesise that such a low explanatory power should be ascribed to the presence of additional confounding factors (e.g., dead biomass, community composition) that have previously proven to interfere with the estimation of grassland diversity (Gholizadeh et al., 2019; Rossi et al., 2021a; Schweiger et al., 2015; Villoslada et al., 2020). Thus, we encourage future investigations to systematically consider all possible confounding factors when testing the spectral diversity-biodiversity relationship in different types of grasslands. Finally, to define the actual possibilities and technical constraints of the relationship, future research should aim at identifying the optimal trade-off between the spatial and spectral resolutions of the RS data used to assess plant diversity in grasslands while incorporating the temporal variations in the spectral signal. In this framework, comparing spectral signatures over an entire growing season (spatio-temporal spectral diversity) could be critical in estimating plant diversity.

## Acknowledgements

We thank Riccardo Testolin and Piero Zannini for useful insights and suggestions that greatly served this study. We thank Elisa Thouverai for technical support in the use of the rasterdiv R package. MP was supported by IGA grant from the Faculty of Environmental Sciences at the Czech University of Life Sciences Prague (project n. 2022B0003). MT and DR were partially supported by the H2020 Project SHOWCASE (Grant agreement No 862480). This study was funded within the Horizon Europe project EarthBridge (ID: 101079310). Funded by the European Union. Views and opinions expressed are, however, those of the author(s) only and do not necessarily reflect those of the European Union. Neither the European Union nor the granting authority can be held responsible for them.

